# Correction of RBFOX1 deficit rescues Huntington’s disease mis-splicing and pathology

**DOI:** 10.1101/2024.11.06.622223

**Authors:** David Lozano-Muñoz, Ainara Elorza, Lucía Mayor, María Santos-Galindo, Miriam Lucas-Santamaría, Alberto Parras, José J. Lucas

## Abstract

RNA mis-splicing correction therapies have been developed for neurological disorders like spinal muscular atrophy and neuronal ceroid lipofuscinosis. In Huntington’s disease (HD), pathogenic mis-splicing was initially observed in genes linked to neurodegeneration, such as *HTT* itself, *MAPT*, and *TAF1*. Later, genome-wide analyses identified a broader mis-splicing signature in HD brains, involving additional neurodegeneration-related genes. Correcting each mis-spliced gene individually would be unfeasible, highlighting the need to target upstream splicing factors altered in HD. Our previous motif-enrichment analyses of intronic sequences flanking the exons mis-spliced in HD identified RBFOX and U2AF2 as candidate splicing factors, both of which are reduced in HD brains. In this study, we tested their pathogenic relevance generating conditional transgenic mouse models that overexpress RBFOX1 or U2AF2 in forebrain neurons and combining them with HD mice. Our results show that moderate overexpression of RBFOX1, but not U2AF2, corrects multiple HD-associated mis-splicing events and alleviates HD mice neuropathology and motor symptoms. These findings demonstrate that RBFOX1 downregulation contributes to HD pathology and underscore the therapeutic potential of strategies aimed at increasing RBFOX1 levels.

## Introduction

Alternative splicing (AS) of pre-mRNA is the differential processing of introns and exons that results in multiple transcript isoforms from individual genes, thereby increasing diversity of the proteome. However, when it is not correctly executed, AS leads to mis-splicing, and this may result in proteins with altered function and stability (Montes *et al*, 2019). Multiple splicing factors and other RNA-binding proteins (RBPs) are responsible for the precise modulation of AS (Ule & Blencowe, 2019), and mis-splicing has been proposed to play a role in multiple brain pathologies, including neurodegenerative diseases (Nik & Bowman, 2019) like Alzheimer (Hsieh *et al*, 2019; Raj *et al*, 2018), amyotrophic lateral sclerosis (Fratta *et al*, 2018; Luisier *et al*, 2018) or Huntington (Elorza *et al*, 2021; Lin *et al*, 2016b; Schilling *et al*, 2019).

Huntington’s disease (HD) is an autosomal dominant neurodegenerative disorder characterized by motor abnormalities together with psychiatric symptoms and by brain atrophy particularly affecting the striatum and the cerebral cortex (Walker, 2007). HD is caused by a CAG repeat expansion in exon 1 of the Huntingtin (*HTT*) gene, encoding a polyglutamine (polyQ) stretch in the N-terminal region of the HTT protein (HDCRG, 1993) with a well-documented pathogenic role (Bates *et al*, 2015). Similar polyQ-encoding CAG triplet repeats expansions cause eight additional neurological disorders, such as some spinocerebellar ataxias (SCAs) and spinal-bulbar muscular atrophy (SBMA) (Orr & Zoghbi, 2007).

Nevertheless, beyond polyQ-driven toxicity, growing evidence indicates that expanded CAG triplet-containing mRNAs cause additional toxicity in these and other triplet repeat disorders such as myotonic dystrophy type 1 (DM1) (Li *et al*, 2008; Ranum & Cooper, 2006). Particularly, it has been suggested an aberrant direct interaction of the expanded CAG triplet repeat containing mRNAs with certain splicing factors (SFs) such as MBNL1 (Mykowska *et al*, 2011), U2AF2 (Schilling *et al*., 2019; Tsoi *et al*, 2011) and SRSF6 (Schilling *et al*., 2019). Accordingly, mis-localization of specific SFs in nuclear foci and inclusion bodies has been reported in HD (de Mezer *et al*, 2011; Fernandez-Nogales *et al*, 2014; Mykowska *et al*., 2011).

The detection of alterations in the alternative splicing machinery in HD was followed by the identification of individual mis-splicing events that could significantly contribute to HD pathogenesis, such as an intron retention in *HTT* that favors a highly toxic exon1-encoded form of the protein (Sathasivam *et al*, 2013), the increased inclusion of *MAPT* exon 10 leading to a detrimental increase in four-tubulin binding repeat-tau (Fernandez-Nogales *et al*., 2014) and the use of an alternative splice site in exon 5 of *TAF1* that correlates with decreased expression of this transcription factor (Hernandez *et al*, 2020), as in X-linked dystonia parkinsonism (Diaw & Lohmann, 2020; Makino *et al*, 2007).

Apart from the mentioned individually identified pathogenic mis-splicing events, through a human/mouse intersect RNA-seq analysis, we were able to define a genome-wide striatal mis-splicing signature triggered by the HD-causing mutation in both mice and humans (Elorza *et al*., 2021; Xing *et al*, 2021), which allowed us to identify additional likely effectors in HD pathogenesis. More precisely, we found six neurodegeneration-linked genes whose mis-splicing correlates with their decreased protein levels. These are: *CCDC88C* (linked to SCA40, OMIM #616053), *KCTD17* [linked to myoclonic dystonia 26 (DYT26) OMIM #616398], *SYNJ1* [linked to early onset Parkinson disease 20 (PARK20), OMIM #615530], *VPS13C* [linked to early onset Parkinson disease 23 (PARK23), OMIM #616840], *TRPM7* [linked to amyotrophic lateral sclerosis-parkinsonism/dementia complex (ALSP/DC), OMIM #105500] and *SLC9A5* [which has been suggested to be linked to episodic kinesigenic dyskinesia 2 (EKD2)](Elorza *et al*., 2021).

Correction of each of the above mentioned potentially pathogenic mis-splicing events might be therapeutically relevant, but approaching them individually would be inefficient and unaffordable. Alternatively, the identification of pivotal upstream splicing factors altered in HD may lead to design therapeutic strategies to simultaneously amend multiple neurodegeneration-associated mis-spliced genes. In this regard, in our previous study we identified five families of SFs that are good candidates to underlie the HD-associated mis-splicing signature because they show decreased protein levels in brains of HD patients and mice, and their consensus sequences are overrepresented in the intronic regions flanking the exons that are mis-spliced in HD (Elorza *et al*., 2021). Out of These five families of SFs, the RBFOX family binds the (U)GCAUG consensus sequence while the rest; namely ELAVL, HNRNPC, TIA1 and U2AF2; bind U-rich motifs.

The RBFOX family of SFs includes RBFOX1 (also known as Ataxin-2 binding protein 1, A2BP1), which is expressed selectively in neurons and in heart and skeletal muscle; RBFOX2 (RBM9), which has a wider expression pattern (whole embryo, ovary, stem cells, brain and skeletal muscle); and RBFOX3 (NeuN), which is expressed exclusively in neurons (Kim *et al*, 2009; Kuroyanagi, 2009). Remarkably, the protein levels of these three splicing factors are decreased in brains of HD patients and mice (Elorza *et al*., 2021).

Out of the U-rich motif binding candidate SFs, U2AF2 (also called U2AF65) is particularly interesting because, apart from being an essential SF (as it is required for the binding of U2 snRNP to the pre-mRNA branch site), it also participates in alternative splicing (Shao *et al*, 2014) and has previously been related to HD pathophysiology (Sun *et al*, 2015; Tsoi *et al*., 2011). Furthermore, a functional interactor of U2AF2 is RBM5 (Bechara *et al*, 2013; Zhou *et al*, 2018), a splicing regulator that displays a differential RNA-binding profile in the R6/2 mouse model and contributes to mis-splicing of neurodegeneration related genes (Mullari *et al*, 2023).

We therefore reasoned that correcting the decrease of either RBFOX or U2AF2 might have a pleiotropic beneficial effect for HD by simultaneously amending multiple mis-spliced genes, and that this could be tested through a mouse genetics approach. For this, here we have generated two new transgenic mouse lines, one overexpressing RBFOX1 and another overexpressing U2AF2, to be combined with HD mouse models in order to explore whether counteracting the decreased levels of these SFs results in attenuation of the molecular, histological and behavioral phenotypes of HD mice.

## Results

### Generation of Tg mouse lines with conditional neuronal overexpression of RBFOX1 or U2AF2

In order to generate RBFOX overexpressing mice, we first considered which RBFOX paralogue would be more relevant to the mis-splicing signature of HD striatum. Interestingly, our previous data from RNA-seq analysis of the widely used R6/1 mouse model of HD at early symptomatic stages of disease (3.5 month-old) showed down-regulation of *Rbfox1* mRNA (FC = 0.8; *Padj =* 2,96 x 10^-5^) (Elorza *et al*., 2021), while *Rbfox2* and *Rbfox3* did not differ between WT and R6/1 mice (Fig. EV1A). Furthermore, we here corroborate the decrease in *Rbfox1* transcript levels by RT-PCR in an independent set of samples from early symptomatic R6/1 mice (25.8% decrease, *P* = 0.03) (Fig. EV1B). This points to Rbfox1 as the earliest altered and therefore best candidate to, upon overexpression, correct mis-splicing and pathology in HD mice. Regarding the different RBFOX1 isoforms, we chose to overexpress the RBFOX1 transcript isoform that lacks the alternatively spliced exon A53, as this preserves the nuclear localization signal required for RBFOX1 to act as a SF (Damianov & Black, 2010; Lee *et al*, 2009; Nakahata & Kawamoto, 2005), as opposed to the cytoplasmic forms of RBFOX1 involved in mRNA stability and translation (Conboy, 2017). Furthermore, we verified by immunohistochemistry staining in striatal sections from R6/1 mice and HD patients that the decrease in Rbfox1 protein previously observed by Western blot, predominantly corresponds to nuclear forms (Fig. EV1C,D). To generate U2AF2 overexpressing mice, no equivalent considerations apply, and we simply decided to express the transcript encoding the 475 aa canonical isoform (Uniprot P26368-1).

Given that mutations in RBFOX or U2AF2 lead to neurodevelopmental disorders (Bill *et al*, 2013; Li *et al*, 2024), to avoid confounding neurodevelopmental artifacts, we decided to use the double transgenic tTA (Tet-Off) system that allows to generate mice with different time windows and levels of transgene expression. With this purpose, we generated β-Gal-BiTetO-hRBFOX1 and β-Gal-BiTetO-hU2AF2 transgenic mouse lines, harboring a bidirectional tetracycline-controlled promoter (BitetO) flanked by the hRBFOX1 or hU2AF2 sequence on one direction, and the β-Gal sequence on the other, the latter as a reporter of transgene expression (Fig. 1A).

**Figure 1.**
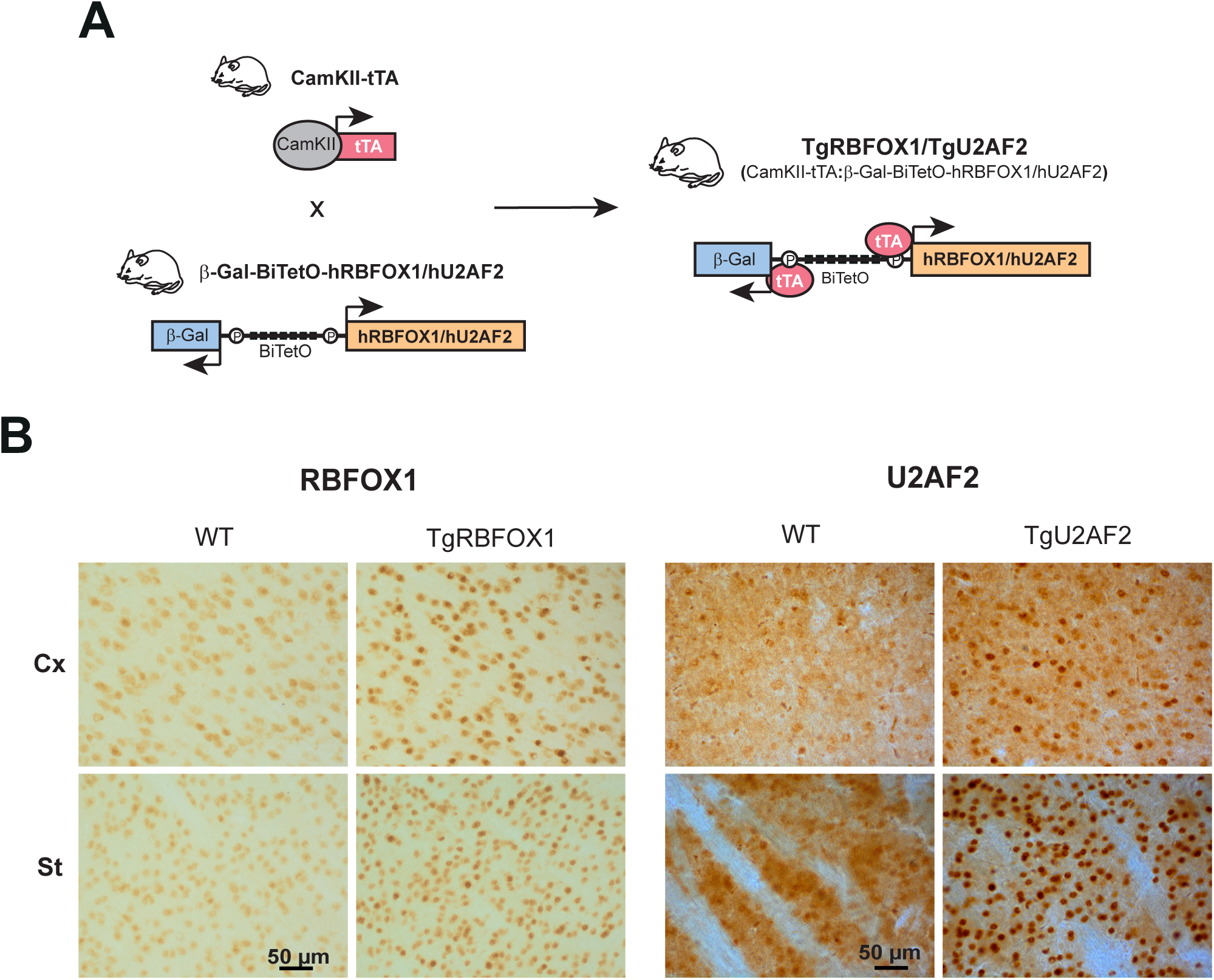
Generation of transgenic mice with conditional overexpression of RBFOX1 nuclear isoform or U2AF2 in forebrain neurons. (**A**) Mice expressing tTA under control of the CamKII promoter (CamKII-tTA mice) were bred with mice carrying the β-Gal-BiTetO-RBFOX1 or β-Gal-BiTetO-U2AF2 construct to yield TgRBFOX1 (CamKII-tTA:β-Gal-BiTetO-RBFOX1) mice or TgU2AF2 (CamKII-tTA:β-Gal-BiTetO-U2AF2) . (**B**) Immunohistochemistry with anti-RBFOX1 or anti-U2AF2 antibody in sagittal sections from 1.5 month-old WT, TgRBFOX1 or TgU2AF2 mice.

The BitetO promoter in β-Gal-BiTetO-hRBFOX1 and β-Gal-BiTetO-hU2AF2 mice is essentially silent and, to achieve transgene transactivation in forebrain regions affected in HD -such as the striatum and the cortex-, we bred these mice with two different CamKII-tTA mouse lines that express the transactivator tTA in forebrain neurons (Fig. 1A). The double transgenic mice are termed TgRBFOX1 or TgU2AF2 and will express the corresponding SF at different levels and time points, according to the specific CamKII-tTA driver mouse line used. The CamKII-tTA mouse line with higher tTA expression (starting at late embryonic stages) resulted in TgRBFOX1 and TgU2AF2 mice with detectable microcephaly (9.4%, *P* = 8 x 10^-3^ and 9.8%, *P* = 1.4 x 10^-3^ decrease in brain weight, respectively), while the TgRBFOX1 and TgU2AF2 mouse lines generated with milder, and only postnatal expression, showed unaltered brain weight (Fig. EV2A,B). For this study aiming correction of phenotypes of HD mice, we therefore decided to use the TgRBFOX1 and TgU2AF2 mice generated with the CamKII-tTA mouse line with moderate level of expression to be combined with HD mice. These mice showed the expected neuronal pattern of transgene expression restricted to forebrain regions such as cortex, striatum and hippocampus, as evidenced by the reporter β-Gal (Fig. EV2C). Furthermore, we verified the nuclear overexpression of RBFOX1 and U2AF2 in the brain regions that undergo the most marked atrophy in HD (cortex and striatum) by immunohistochemistry with antibodies against the corresponding SF (Fig. 1B).

### Overexpression of RBFOX1, but not U2AF2, attenuates the HD-associated motor deficit of R6/1 mice

To check whether correction of the RBFOX1 or U2AF2 deficit can be beneficial to HD mice, we combined R6/1 mice with TgRBFOX1 or TgU2AF2 mice. Then, the resulting WT, R6/1 and R6/1+TgRBFOX1 or WT, R6/1 and R6/1+TgU2AF2 littermates were subjected to the accelerating rotarod test, at ages at which R6/1 mice are known to show early (3.5 months) or advanced (5 months) phenotypes (Fig. 2A). Remarkably, we observed that overexpression of RBFOX1 attenuated the motor phenotype of R6/1 mice. More precisely, at the age of 3.5 months, R6/1+TgRBFOX1 mice performed better than R6/1 mice (Fig. 2B) and they still showed a tendency to perform better than R6/1 mice at the age of 5 months. However, overexpression of U2AF2 did not attenuate the motor phenotype of HD mice, as R6/1+TgU2AF2 mice were indistinguishable from R6/1 mice at both tested ages (Fig. 2B). These results strongly suggest that mis-splicing of RBFOX targets, as a consequence of decreased levels of RBFOX1, significantly contributes to HD pathogenesis.

**Figure 2.**
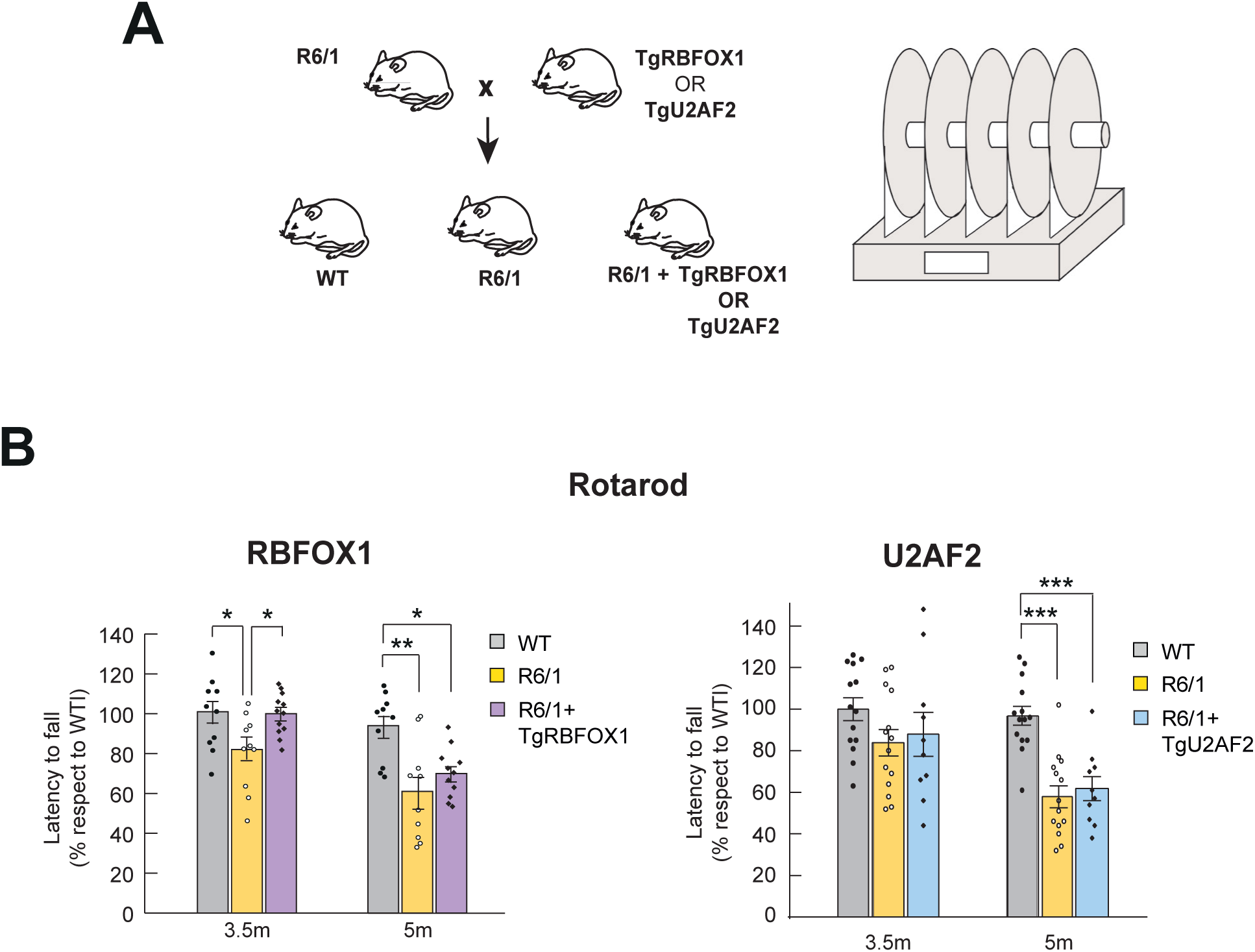
Overexpression of RBFOX1 but not U2AF2 delays HD-associated behaviors in R6/1 mice. (**A**) TgRBFOX1 and TgU2AF2 mice lines were bred with R6/1 mice, yielding R6/1+TgRBFOX1 or R6/1+TgU2AF2 which were tested in accelerating rotarod apparatus. (B) Evolution of the percentage of the mean latency to fall in the accelerating rotarod test of control, R6/1 and TgRBFOX1 or TgU2AF2, mice at different ages. Analysis of variance (ANOVA), followed by Tukeýs post-hoc test. Data represent mean ± SEM. (*P <0.05, ** P < 0.01, *** P < 0.001)

### RBFOX1 correction in HD mice attenuates mis-splicing of disease associated genes

To explore RBFOX1 target transcripts relevant to HD pathophysiology, we performed RNA-seq analysis of forebrain mRNA from TgRBFOX1 mice, for comparison with the mis-splicing signature previously stablished for HD striatum. The splicing events that varied between WT and TgRBFOX1 mice with a difference in percent-splice-in (dPSI) > 10 were identified by running Vast-tools (Fig. 3A). Then, the 332 genes with exons altered in TgRBFOX1 mice were confronted to the 245 genes that we previously found with mis-spliced cassette exons is striatum of both HD patients and early symptomatic R6/1 mice (Elorza *et al*., 2021). We observed a significant overlap (RF: 3.3, *P* < 1.47 x 10^-6^) between genes affected by the in vivo overexpression of RBOXF1 and the HD-mis-splicing signature (approximately 9% of the latter) (Fig. 3B), as expected given the high percentage of direct RBFOX targets -previously identified through CLIP-seq analysis (Weyn-Vanhentenryck *et al*, 2014)-, within the HD mis-splicing signature (Fig. EV3A). Interestingly, out of the six key HD mis-spliced genes previously identified as likely pathogenic (Elorza *et al*., 2021), two (Synj1 and Slc9a5) were altered upon *in vivo* overexpression of RBFOX1 (Fig. 3B).

**Figure 3.**
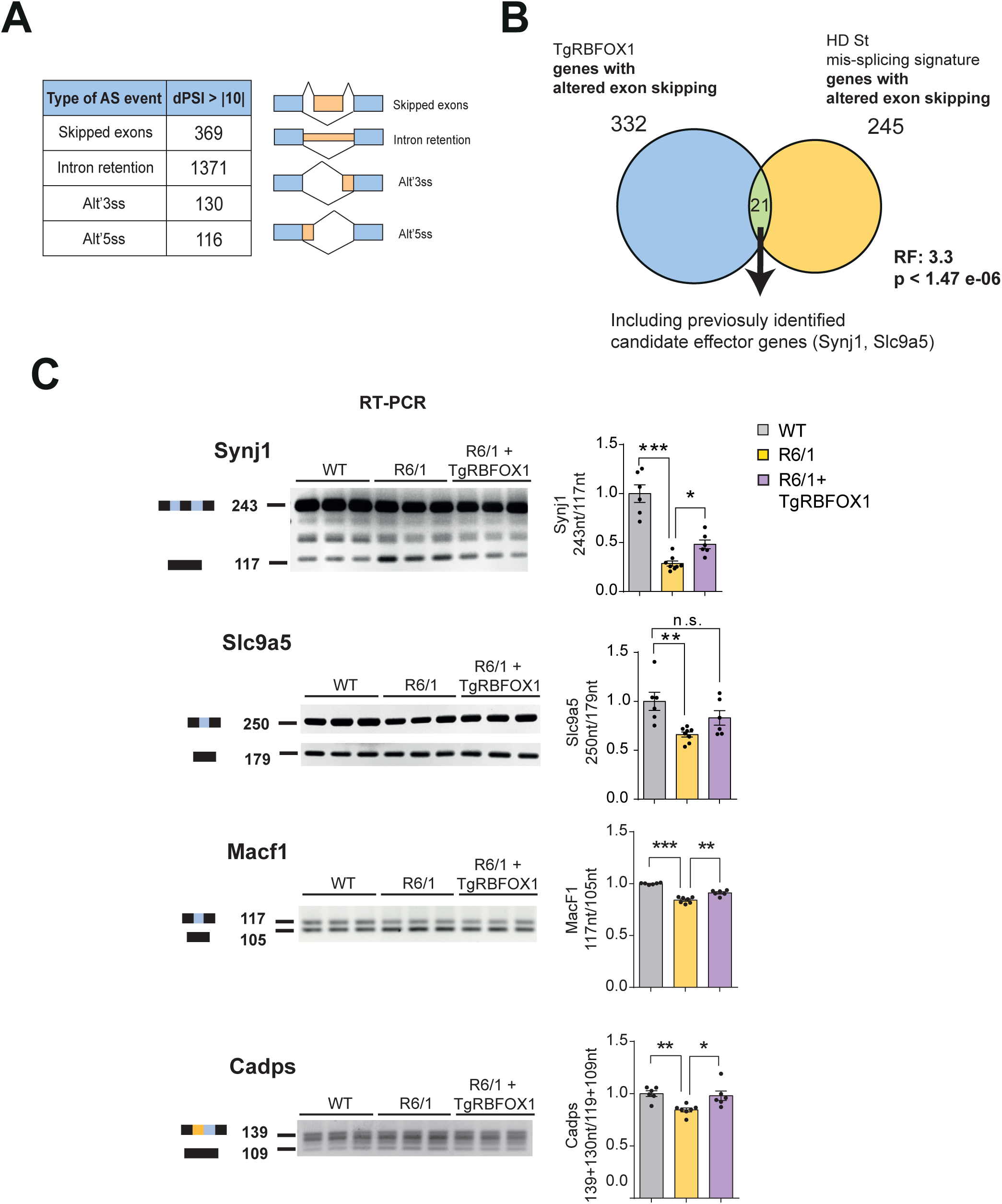
Overexpression of RBFOX1 nuclear isoform attenuates Huntington’s disease-associated mis-splicing in R6/1 mice. RNA-seq was performed on forebrain of WT (n=2) and TgRBFOX1 P5 mice (n=2), and splicing was analyzed using Vast-tools (**A**) The different types of alternative splicing events with an absolute dPSI > 10 between wild-type and TgRBFOX1 mice (**B**) Venn diagram showing the 21 genes in the intersection between the 245 genes with exons mis-spliced in Huntington’s disease^15^ and the 332 genes with exons with changes in splicing in TgRBFOX1. (**C**) Representative RT-PCR image and quantification of mis-spliced events in cortex of 3.5-month-old WT (n = 6), R6/1 (n = 7) and R6/1+TgRBFOX1 (n = 6) mice (ANOVA followed by Tukey’s or Games-Howell post hoc test; *P < 0.05; **P < 0.01; ***P < 0.001). Graphs show means ± SEM.

We then decided to validate by RT-PCR the correction in R6/1+TgRBFOX1 mice of the mis-splicing of Synj1 and Slc9a5 that takes place in R6/1 mice, as well as that other disease-associted genes that are direct RBFOX-targets and have exons mis-spliced in brains of HD patients and mice (such as Macf1 and Cadps). As shown in Fig.3C, the mis-splicing of Synj1, Macf1 and Cadps was total or partially corrected in R6/1+TgRBFOX1 mice. Regarding Slc9a5, we noticed that despite not being included in the set of direct RBFOX targets of Fig. 3A, it contains the (U)GCAUG RBFOX binding consensus sequence in the intron downstream the 71 nt exon whose inclusion is decreased in R6/1 mice compared to WT mice and, interestingly, such decreased inclusion is no longer observed in R6/1+TgRBFOX1 mice (Fig. 3C). In summary, from these experiments we can conclude that mild RBFOX1 overexpression in forebrain neurons attenuates the mis-splicing of disease associated genes that takes place in HD mice.

### RBFOX1 correction attenuates neuropathology in HD mice

We finally decided to check whether partial correction of the Rbfox1 decrease of R6/1 mice together with its subsequent attenuation of mis-splicing of RBFOX targets, apart from attenuating motor phenotype, also has a positive impact on HD-associated neuropathology. For this, we first immunostained sagittal sections from 3.5-month-old WT, R6/1 and R6/1+TgRBFOX1 mice for the striatal marker DARPP32 to analyse atrophy (Fig. 4A) and for cleaved caspase 3 to detect apoptotic cells (Fig. 4B). This revealed that the decrease in striatal area observed in R6/1 mice respect to WT mice was significantly attenuated in R6/1+TgRBFOX1 mice. In good agreement, the significant increase in the number of cleaved caspase-3 positive cells detected in R6/1 mice is no longer observed in R6/1+TgRBFOX1 mice. Altogether, our results demonstrate that mild transgenic neuronal overexpression of RBFOX1 suffices to attenuate the mis-splicing and the neuroanatomical and motor abnormalities of HD mice, thus evidencing that the decrease of RBFOX1 that takes place in brains of HD patients and mice significantly contributes to HD pathogenesis.

**Figure 4.**
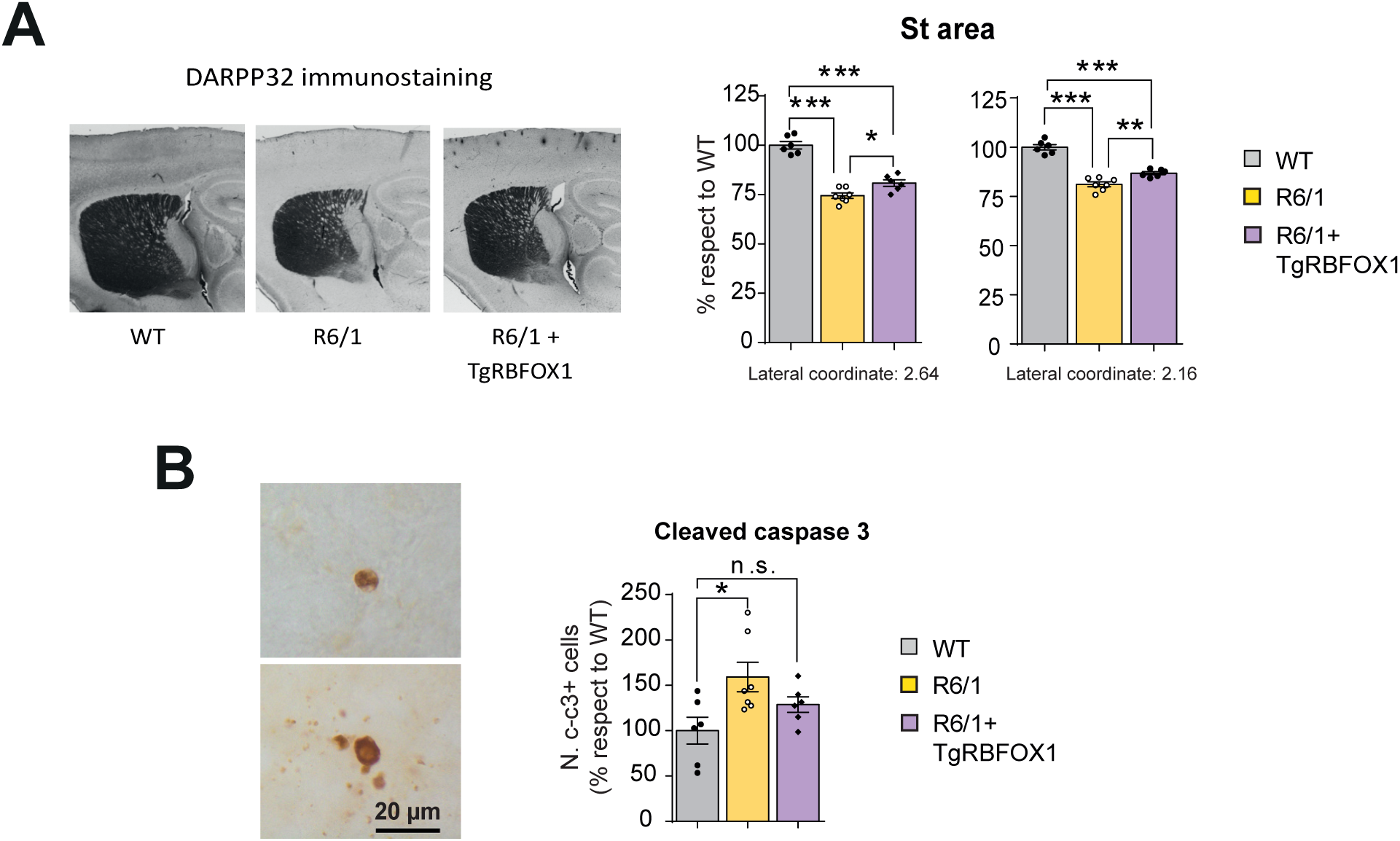
Overexpression of RBFOX1 nuclear isoform attenuates striatal atrophy in R6/1 mice. (**A**) Representative images of DARPP32-immunostained striatal (St) area and its quantification in sagittal sections at two different lateral coordinates of the mouse brain atlas for WT (n = 6), R6/1 (n = 7) and R6/1+TgRBFOX1 (n = 6) mice at 3.5 months of age (ANOVA, followed by Tukey’s post hoc test; *P < 0.05; ***P < 0.001) (**B**) Representative images showing immunostaining of cleaved caspase-3 (left) and quantification of cleaved caspase-3-positive cells in WT (n = 6), R6/1 (n = 7) and R6/1+TgRBFOX1 (n = 6) mice (right) (ANOVA, followed by Tukey’s post hoc test; \**P* < 0.05; n.s., not significant).

## Discussion

Here we generate transgenic mouse lines with neuronal overexpression of RBFOX1 or U2AF2 to attenuate the decreased levels of these splicing factors in HD mice. The observed amelioration of transcriptomic, histopathological, and behavioral phenotypes upon RBFOX1 overexpression demonstrates that diminished RBFOX activity contributes to HD pathogenesis, and suggests therapeutic potential of interventions able to increase RBFOX activity.

RBFOX1 was one of the candidate SFs to overexpress in HD mice because it is the first member of the RBFOX family to show decreased transcript and protein levels. However, along disease progression, the three members of the family show decreased protein levels. Understanding the mechanisms by which their levels decrease may help to envision ways to prevent such decrease. As mentioned, early decrease of RBFOX1 protein levels can be explained by its also decreased transcript levels, but early decreases of RBFOX2 and RBFOX3 do not correlate with diminished transcript levels, suggesting that post-transcriptional mechanisms might also be at play. In this regard, CPEB4-dependent alterations in transcriptome polyadenylation and subsequent translation of affected transcripts has been reported in HD mice and, interestingly, Rbfox2 transcript is among those showing reduced polyadenylation in R6/1 mice (Picó *et al*, 2021), what is expected to contribute to decreased translation (Ivshina *et al*, 2014).

Regulation of transcript stability and translation by microRNAs (miRNAs) is another mechanism of posttranscriptional regulation of gene expression that is known to affect RBFOX genes. At least two miRNAs have been shown to diminish RBFOX1 expression: miR-980 in the context of stress-dependent ribonucleoprotein granule formation (Kucherenko & Shcherbata, 2018) and miR-129-5p that promotes synaptic downscaling in models of chronically elevated network activity (Rajman *et al*, 2017). In this regard, hyperactivity of striatal neurons has been reported in HD mice (Miller *et al*, 2011; Rebec *et al*, 2006), and one might expect a compensatory increase of mir-129-5p. However, no changes in mir-129 levels have been reported in the striatum of individuals with HD, while in the frontal cortex they show upregulated mir-129-3p and downregulated mir-129-5p (Marti *et al*, 2010).

In any case, neutralizing miR-980 or miR-129-5p might have the desirable effect of increasing RBFOX1 protein levels. Antagomirs are a class of chemically engineered oligonucleotides designed to silence endogenous microRNAs as their sequence is complementary to their specific miRNA target (Stenvang *et al*, 2012). Interestingly, antagomirs have already been shown to be effective and safe in animal models of neuromuscular diseases such as DM1 upon systemic administration (Cerro-Herreros *et al*, 2018). In that study, the authors sought to increase levels of a different splicing factor (MBNL) by antagonizing miR-23b and miR-218, and the target tissue was skeletal muscle. However, reaching the brain to increase RBFOX1 expression with potential miR-980 or miR-129-5p antagomirs can be more challenging as they should be engineered to cross the blood-brain barrier. It is worth noting though, that we have limited our study to analyzing RBFOX levels and functional implications in the brain of HD mice and patients but, since RBFOX-dependent AS is also important in muscle physiology (Singh *et al*, 2018) and this tissue is also affected in HD (Zielonka *et al*, 2014), we cannot fully discard a role of RBFOX also in HD peripheral tissues. However, exploring this goes beyond the scope of the present study.

As mentioned, RBFOX1 loss of function, normally due to deletions or point mutations in the *RBFOX1* locus, has previously been associated to neurodevelopmental disorders like epilepsy, intellectual disability or ASD (Bhalla *et al*, 2004; Bill *et al*., 2013; Lim *et al*, 2006; Martin *et al*, 2007; Sebat *et al*, 2007). The relevance of RBFOX1 to ASD pathogenesis is believed to be due, in part, to its ability to regulate microexons (Li *et al*, 2015). Microexons, are exons of 30 or fewer nucleotides with a neural specific alternative splicing (AS) pattern that has been shown to be dysregulated in ASD and to contribute to its aetiology (Gonatopoulos-Pournatzis & Blencowe, 2020; Irimia *et al*, 2014; Li *et al*., 2015; Parras *et al*, 2018). Curiously, two of the four HD-associated mis-splicing events that we show corrected in this study upon transgenic RBFOX1 overexpression, Macf1 and Cadps, are microexons.

Interestingly, a recent study performed on an established isogenic HD cell model demonstrates widespread neuronal differentiation stage- and CAG length-dependent splicing changes (Tano *et al*, 2023) and we noticed that the genes whose mis-splicing gets corrected by our RBFOX1 overexpression approach (Synj1, Macf, Cadps and Slc19a5) are also found mis-spliced in the isogenic HD cell model, thus suggesting that many of the RBFOX1 dependent mis-splicing events might be taking place at early stages of neuronal maturation and prodromal disease. This fits with the mentioned well stablished role of RBFOX1 during neurodevelopment and the neurodevelopmental component of HD pathogenesis (Braz *et al*, 2022).

Interestingly, it has also been reported an increase in RBFOX1 levels and an associated aberrant pattern of alternative RNA-processing in neurons derived from iPSCs of Parkinsońs disease (PD) patients (Lin *et al*, 2016a), In this regard, our results show that excess RBFOX1 in our mouse lines with highest transgene expression results *per se* in decreased brain weight. Besides, in combination with R6/1 mice, these mice -far from alleviating the motor impairment of HD mice-, even seem to induce a worsening (data not shown). Accordingly, if the here generated TgRBFOX1 mouse lines were bred with proper tTA driver mice to drive RBFOX1 overexpression to dopaminergic neurons, they might represent a model of excess RBFOX activity in PD.

Although our previous scan motif analysis identified U2AF2 as a possible upstream splicing factor responsible of the HD mis-splicing signature, and two previous studies had related U2AF2 to HD pathophysiology (Sun *et al*., 2015; Tsoi *et al*., 2011), the results of this study indicate that the decrease in U2AF2 levels, at least in striatal and cortical neurons, does not seem to significantly contribute to disease progression. The possibility still exists that overexpressing U2AF2 in other cell types where expression of mutant Htt is also pathogenic, like astrocytes (Bradford *et al*, 2009) or oligodendrocytes (Ferrari Bardile *et al*, 2019) might have resulted in amelioration of HD-like phenotypes. These seems however unlikely, as gene ontology analyses of genes accounting for the HD mis-splicing signature essentially yielded neuronal terms. An alternative explanation for the lack of improvement of HD mice upon U2AF2 overexpression is that, as mentioned, in our previous motif analysis (Elorza *et al*., 2021) we also identified other U-rich binding SFs, such as PTBP-1/2 or HNRNPC. As these U-rich motifs are highly similar, it is challenging to specifically distinguish each signal. Moreover, previous studies have shown that these splicing factors and U2AF2 compete at various splice sites (Sutandy *et al*, 2018; Zarnack *et al*, 2013). Therefore, it is possible that the U2AF2 signal detected in our previous analysis could be attributed to other splicing factors.

In summary, here we demonstrate that RBFOX1 plays a critical role upstream of the mis-splicing signature of HD brains, as a moderate increase of RBFOX1 in forebrain neurons by genetic manipulation of HD mice attenuates mis-splicing and ameliorates their neuropathology and motor symptoms. This study therefore demonstrates that RBFOX1 contributes to HD pathogenesis and suggests that RBFOX1 increasing strategies may represent a new avenue for therapeutic intervention for HD.

## Methods

### Human brain tissue samples

Brain specimens used in this study from HD patients and controls (CTRL) were provided by Institute of Neuropathology Brain Bank (HUB-ICO-IDIBELL, Hospitalet de Llobregat, Spain), the Neurological Tissue Bank of the IDIBAPS Biobank (Barcelona, Spain), the Banco de Tejidos Fundación Cien (BT-CIEN, Madrid, Spain) and the Netherlands Brain Bank (Amsterdam, The Netherlands). Written informed consent for brain removal after death for diagnostic and research purposes was obtained from brain donors and/or their next of kin. Procedures, information and consent forms were reviewed and approved by the Bioethics Subcommittee of Consejo Superior de Investigaciones Científicas (CSIC, Madrid, Spain).

### Mice

R6/1 transgenic mice for the human exon-1-Htt gene (Mangiarini *et al*, 1996) were maintained in B6CBAF1 background; CamKII-tTA mouse lines (Mayford *et al*, 1996) were maintained in a pure C57BL/6J background. Mice with transgenic expression of human RBFOX1 or human U2AF2 under a tetracycline-regulated promoter (TgRBFOX1 or TgU2AF2) were generated for this study (for details see ‘Generation of TgRBFOX1 and TgU2AF2 mice’ below) and were maintained in C57BL/6J background. All animals were bred and housed at the Centro de Biología Molecular Severo Ochoa animal facility. Mice were housed four per cage with food and water available *ad libitum* and maintained in a temperature-controlled environment on a 12/12 h light-dark cycle with light onset at 08:00. Animal housing and maintenance protocols followed the guidelines of Council of Europe Convention ETS123. Animal experiments were performed under Comunidad de Madrid Regional Government protocols (PROEX 293/15 and PROEX 247.1/20) and approved by CSIC’s Ethics Committee and Centro de Biología Molecular Severo Ochoa’s Committee for Animal Care and Utilization (Comité de Ética de Experimentación Animal del CBM, CEEA-CBM), Madrid, Spain.

### Generation of RBFOX1 and U2AF2 transgenic mice

Human RBFOX1 cDNA (from pENTR-A2BP1, Addgene, 16176) or human U2AF2 cDNA (from pCDNA3-U2AF65-DYK, GenScript, OHu17874) were independently cloned into a plasmid containing a bidirectional TetO sequence that also harbors the β-galactosidase (β-Gal) reporter with a NLS (pBI-G, Clontech, 631004). The resulting construct was linearized and microinjected into single-cell C57BL/6JxCBA embryos. This resulted in β-Gal-BiTetO-hRBFOX1 or β-Gal-BiTetO-hU2AF2 founder mice that were backcrossed into a pure C57BL/6J background. These mice were then crossed with CamKII-tTA mice in C57BL/6J background to obtain conditional double transgenic mice with forebrain neuronal expression of hRBFOX1 or hU2AF2 (TgRBFOX1 and TgU2AF2 mice).

### Rotarod test

Motor coordination was assessed with an accelerating rotarod apparatus (Ugo Basile, Comerio, Italy). Mice were trained during two consecutive days, the first day with four trials at fixed 4 rpm for 1-min each and the second day with four trials of 2 min (the first minute at 4 rpm and the second minute at 8 rpm). On the third day, rotarod was set to accelerate from 4 to 40 rpm over 5 min and mice were tested in four trials. The latency to fall from the rotarod was measured as a mean of the four accelerating trials.

### Tissue preparation for staining

For human samples, formalin-fixed (4%, 24 h), paraffin-embedded tissue from striatum were used. Sections (5-μm thick) were mounted on superfrost-plus tissue slides (Menzel-Gläser) and deparaffinized. For mouse samples, mice were killed using CO2. Brains were quickly extracted and the left hemisphere was immersed in 4% paraformaldehyde overnight. After profuse washing in PBS, hemispheres were immersed in sucrose 30% in PBS for at least 72 h and then included in OCT (Optimum Cutting Temperature compound, Tissue-Tek, Sakura Finetek Europe, ref. 4583), frozen and stored at -80°C until use. Mouse sagittal sections (30 µm thick) were sequentially cut on a cryostat (Thermo Scientific), collected and stored free floating in glycol-containing solution (30% glycerol, 30% ethylene glycol in 0.02M phosphate buffer) at -20°C.

### Immunohistochemistry

For human samples, peroxidase activity was quenched with 0.3% H2O2 in methanol for 30 min, followed by antigen retrieval with 10 mM pH 6.0 citrate buffer heated in microwave for 15 min. For mouse sections, tissues were first washed in PBS and then immersed in 0.3% H2O2 in PBS for 45 min to quench endogenous peroxidase activity. After PBS-washes, sections were immersed for 1 h in blocking solution [PBS containing 0.5% foetal bovine serum, 0.3% Triton X-100 and 1% bovine serum albumin] and incubated overnight at 4 °C with primary antibody diluted in blocking solution. After washing, brain sections were incubated first with biotinylated goat anti-rabbit or anti-mouse secondary antibody and then with avidin-biotin complex using the Elite Vectastain kit (Vector Laboratories, PK-6101 and PK-6102). Chromogen reactions were performed with diaminobenzidine (SIGMAFAST DAB, Sigma, D4293) for 10 min. Mouse sections were mounted on glass slides and coverslipped with Mowiol (Calbiochem, Cat. 475904), while human sections where first dehydrated and then mounted with DePex (SERVA). Images were captured using an Olympus BX41 microscope with an Olympus camera DP-70 (Olympus Denmark A/S). Antibodies: rabbit anti-RBFOX1 (1:5000, Novus, NBP1-90304), rabbit anti-β-Gal (1:2000, Invitrogen, A-11132), rabbit DARPP32 (1:3000, BD, 611520) and rabbit Cleaved caspase-3 (1:100, Cell Signaling, 9661) for mouse samples. Mouse anti-RBFOX1 (1:5000, Merk Millipore, MABE985) for human samples.

### Quantification of cleaved caspase-3-positive cells

The total number of immunopositive cells in three sagittal sections (lateral coordinates 1.20, 1.44 and 1.68 mm) was quantified for each animal using an Olympus BX41 microscope with an Olympus camera DP-70 (Olympus Denmark A/S). Mean values per genotype were used for statistical comparison.

### Striatal area measurement

The striatal area was measured by Darpp32-positive immunostaining in sagittal sections (1 section/animal) (lateral coordinate 2.64 and 2.16). Images were captured at 2.5x magnification (Canon EOS 450D digital camera) and area quantification was made using ImageJ software. Mean values for each genotype were used for statistical comparison

### RNA sequencing and analysis

Mice were killed using CO2 and mouse brains were quickly dissected on an ice-cold plate and the different structures stored at -80°C. Total RNA from forebrain of P5 WT (n=2) and TgRBFOX1 (n=2) mice was isolated using the Maxwell® 16 RSC simplyRNA Tissue Kit (Promega, AS1340). Total RNA was quantified by Qubit® RNA BR Assay kit (Thermo Fisher Scientific) and the RNA Quality number (RQN) was estimated using the Qsep100TM (BiOptic) detecting, on average, a RQN value over 9. Stranded poly(A) enriched libraries were generated and sequenced on a Novaseq X Plus (Novogene) with a read length of 2 x 150 bp, generating at least 100 million reads per sample.

Vertebrate Alternative Splicing and Transcription Tools (VAST-TOOLS) v2.5.1 was used to analyze AS (Irimia *et al*., 2014). Briefly, reads in FASTQ format were aligned to the mouse junction library (mm10; vastdb.mm2.23.06.20) using align module, and the generated files were joined using the combine module. For differential splicing analysis the module compare was employed with the following parameters: --min_dPSI 10 and –min_range 5.

### Reverse transcriptase quantitative and semiquantitative PCR

Retrotranscription reactions were performed using iScript™ Reverse Transcription Supermix for RT-qPCR (Bio-Rad). Relative quantification was carried out for mRNA analysis in striatum of mice (WT and R6/1). qRT-PCR (CFX 384 Biorad) was carried out with 5 ng of cDNA in a volume of 4 μl with 1 μl of 5 μM forward and reverse primers mix and 5 μl of SsoFast EvaGreen Supermix premix (Biorad, CN172-5204). Triplicate reactions were carried out for each mRNA. The following amplification protocol was used: initial denaturation of 5 s at 95 °C + 40 cycles x [5s at 95 °C + 5 s at 60 °C] + [5 s at 60 °C + 5 s at 95 °C]. Fluorescence was taken at the end of elongation step. Data were analysed by GenEx 5.3.7 software (Multid AnaLyses AB). The mRNA levels were normalized first relative to total RNA and then relative to the 18S ribosome subunit, β-ACTIN, GAPDH and β-TUBULIN gene expression in each sample.

RBFOX1 transgene and selected events were evaluated with semiquantitative reverse transcription-PCR. cDNA (50 ng) was amplified with specific primers. PCR products were resolved on 2% resolution Metaphor agarose gels (Lonza).

### Primers

Mouse primers use in this study are as follows: Rbfox1 Forward 5′-GACCCCTACCACCACACACT - 3′, Reverse 5′-TCTTGGCATCGGTCAAGG -3′; 18s Forward 5′-CTCAACACGGGAAACCTCAC-3′, Reverse 5′-CGCTCCACCAACTAAGAACG-3′; β-actin Forward 5′-CTAAGGCCAACCGTGAAAAG-3′, Reverse 5′-ACCAGAGGCATACAGGGACA-3′; Gapdh Forward 5′-CTCCCACTCTTCCACCTTCG-3′, Reverse 5′-CATACCAGGAAATGAGCTTGACAA-3′; β-Tubulin Forward 5′-GACCTATCATGGGGACAGTG-3′, Reverse 5′-CGGCTCTGGGAACATAGTTT-3′; Synj1 Forward 5′-CCCAGACTCTAGAGCCCAAGA-3′, Reverse 5′-GCTTGAGGGGAAGGCTGATTAC-3′; MacF1 Forward 5′-CAGCAGGTGTGGCTGTTAGC-3′, Reverse 5′-CCAGACATCAAAGTCAAAGTTGGC-3′; Cadps Forward 5′-ACTGCAACAAAACGAGGAGCA-3′, Reverse 5′-TGCATCCATGTCCACTGCAAA-3′; Slc9a5 Forward 5′-GCTGAGGGTGAAGAGGAGTGA-3′, Reverse 5′-GCTGATGGCATCTCGGATGTT-3′.

Human primers use in this study are as follows: RBFOX1 Forward 5′-GCCACAGCACGTGTAATGACA-3′, Reverse 5′-CCACAACTGGATTCAATTTCCAG-3′; Transgenic RBFOX1 Forward 5′-CCGAAGCGTAAGGCTGAG-3′, Reverse 5′-CCCGAATTCCTGCATCAAG-3′; 18S Forward 5′-ATCCATTGGAGGGCAAGTC-3′, Reverse 5′-GCTCCCAAGATCCAACTACG-3′; β-TUBULIN Forward 5′-CTTTGTGGAATGGATCCCCA-3′, Reverse 5′-GACTGCCATCTTGAGGCCA-3′. *GAPDH* and *β-ACTIN* (TATAA Biocenter, qA-01-0101S, qA-01-0104S)

### Brain weight

Whole brains were extracted from skulls of 1.5 month-old mice and weighted in a precision scale (Mettler Toledo, AB265-S).

### Statistics

Statistical analysis was performed with SPSS 26.0 (SPSS® Statistic IBM®). Data are represented as Mean ± SEM (Standard Error of the Mean). The normality of the data was analysed by Shapiro-Wilk. Homogeneity of variances was analysed by Levene test. For two independent group comparison, two-tailed Student’s t-test (data with normal distribution) or Mann-Whitney U-test (data with non-normal distribution) was performed. For multiple comparisons, data were analysed by one-way ANOVA followed by a Tukey’s post-hoc test (data with normal distribution) or by Games-Howell post hoc test (data with non-normal distribution). A critical value for significance of *P* < 0.05 was used throughout the study. Regarding overlap between HD mis-spliced genes and RBFOX direct and functional targets, enrichment tests were carried out with two-sided Fisher’s exact test.

## Data availability

Data are available from the corresponding author upon reasonable request.

## Acknowledgements

Human tissue was obtained from Institute of Neuropathology (HUB-ICO-IDIBELL) Brain Bank, the Neurological Tissue Bank of the IDIBAPS Biobank, the Banco de Tejidos Fundación CIEN (BT-CIEN, Madrid, Spain), and the Netherlands Brain Bank (Amsterdam, The Netherlands). We also thank excellent technical assistance by Miriam Lucas and by the following core facilities: CBMSO-Genomics & NGS, CBMSO-Animal Facility and allocation of computing time at the Centro de Computación Científica-UAM (CCC-UAM).

This work was supported by CIBERNED - Health Institute Carlos III (ISCIII) Collaborative Grants No. PI2015-2/06-3 and PI2018/06-1; by grants: SAF2015-65371-R (Spanish Ministry of Economy and Competitiveness (MINECO)), RTI2018-096322-B-I00 (Spanish Ministry of Science, Innovation and Universities (MICIU)) and PID2021-123141OB-I00 (AEI, Agencia Estatal de Investigación: Spanish Ministry of Science and Innovation and the European Union) to JJL. Fellowship for doctoral thesis PRE2019-090829 (Spanish Ministry of Science and Innovation) to DL-M. The Center for Molecular Biology Severo Ochoa is a Severo Ochoa Center of Excellence (MICIN, Award CEX2021-001154-S).

## Author contribution

AE, DL-M, LM, MS-G and ML-S performed experiments. AE and DL-M analyzed the data. DL-M and MS-G performed bioinformatics. AP and LM contributed to TgRBFOX1 and TgU2AF2 mice lines generation, respectively. M.M.-S. performed behavioral tests on mice. AE and JJL wrote the paper with input from all authors. The order of authorship has been assigned after discussion and voting by all authors.

## Extended view figure legends

**Figure EV1.**
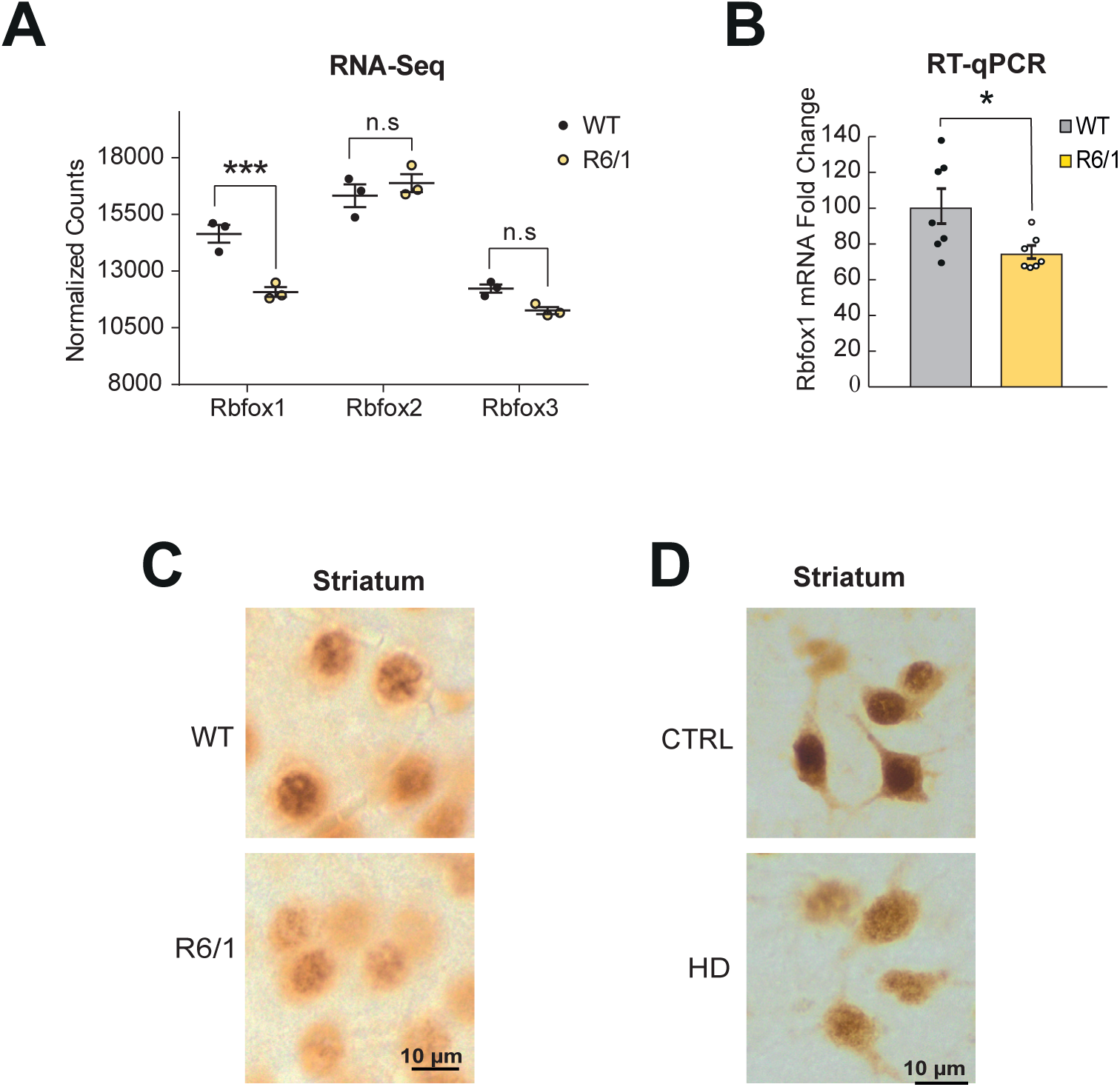
Early decrease of Rbfox1 transcripts in mice and nuclear protein isoforms decrease in Huntington’s disease mice and Huntington’s disease patients. (**A**) Normalized counts of the three RBFOX genes in striatal RNA from 3.5 month-old WT (n=3) and R6/1 mice (n=3) according to RNA-seq datasets in Elorza *et al*.^15^ (**B**) Quantification of *Rbfox1* transcript levels by RT-qPCR in striatal RNA from 3.5 month-old WT (n=7) and R6/1 mice (n=7) (Student’s t-test; \**P* < 0.05). (**C**) Rbfox1 immunohistochemistry in striatum of 3.5 month-old WT and R6/1 mice. (**E**) Representative RBFOX1 immunohistochemistry staining in striatum of control and HD subjects.

**Figure EV2.**
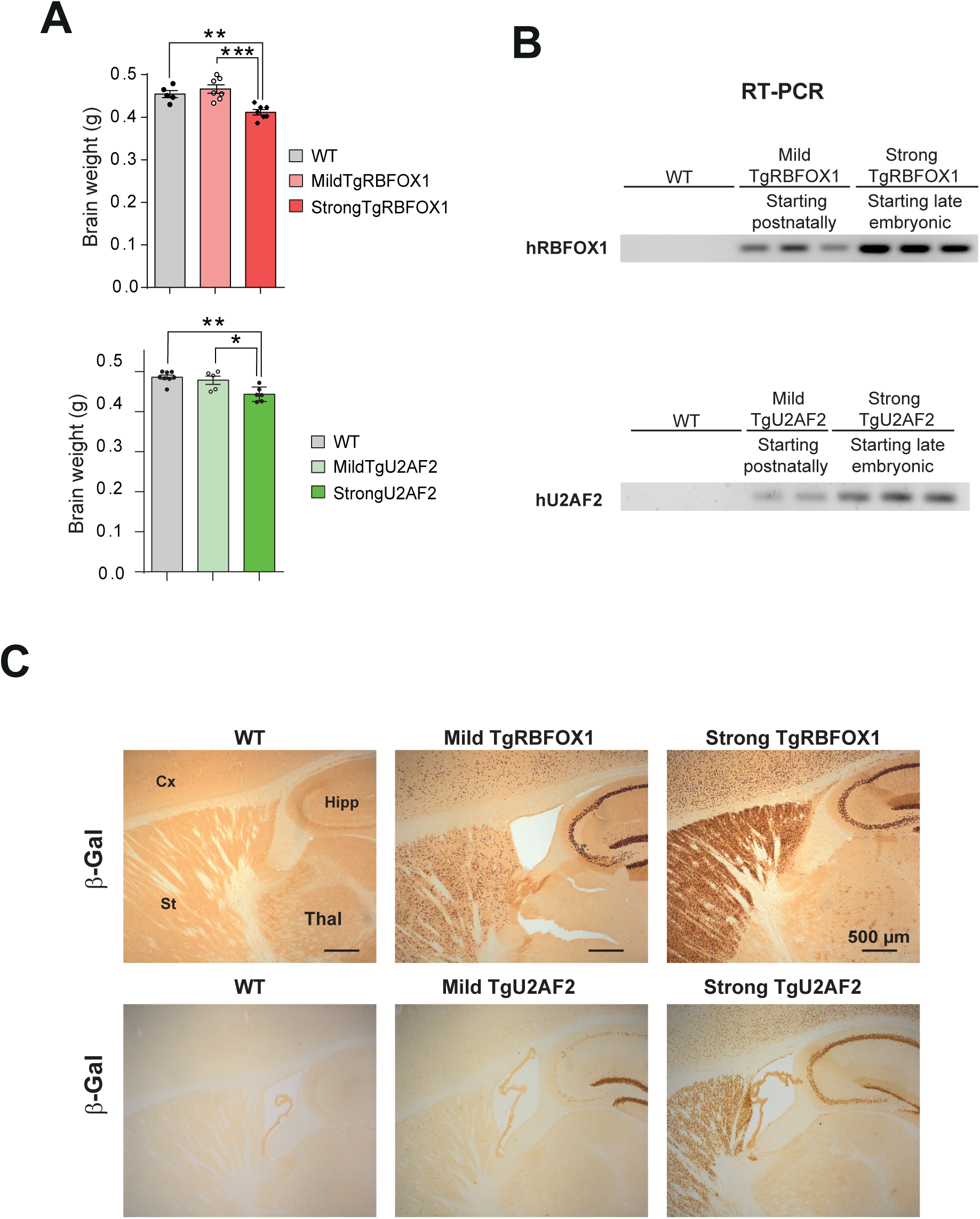
β-Gal-BiTetO-RBFOX1 or β-Gal-BiTetO-U2AF2 bred with CamKII-tTA mouse line with higher tTA expression results in TgRBFOX1 and TgU2AF2 mice with microcephaly. (A) Gel shows RT-PCR amplification of Tg-RBFOX1 or TgU2AF2 mRNA in wild-type, MildTgRBFOX1 or MildTgU2AF2 and StrongTgRBFOX1 or StrongTgU2AF2 mice. (B) Histogram shows brain weight of WT (n=5/n=8), MildTgRBFOX1 (n=7) or MildTgU2AF2 (n=5) and Strong TgRBFOX1 (n=7) or Strong TgU2AF2 (n=6) mice (ANOVA, followed by Tukey’s post hoc test; *P < 0.05;**P < 0.01; ***P < 0.001).

**Figure EV3.**
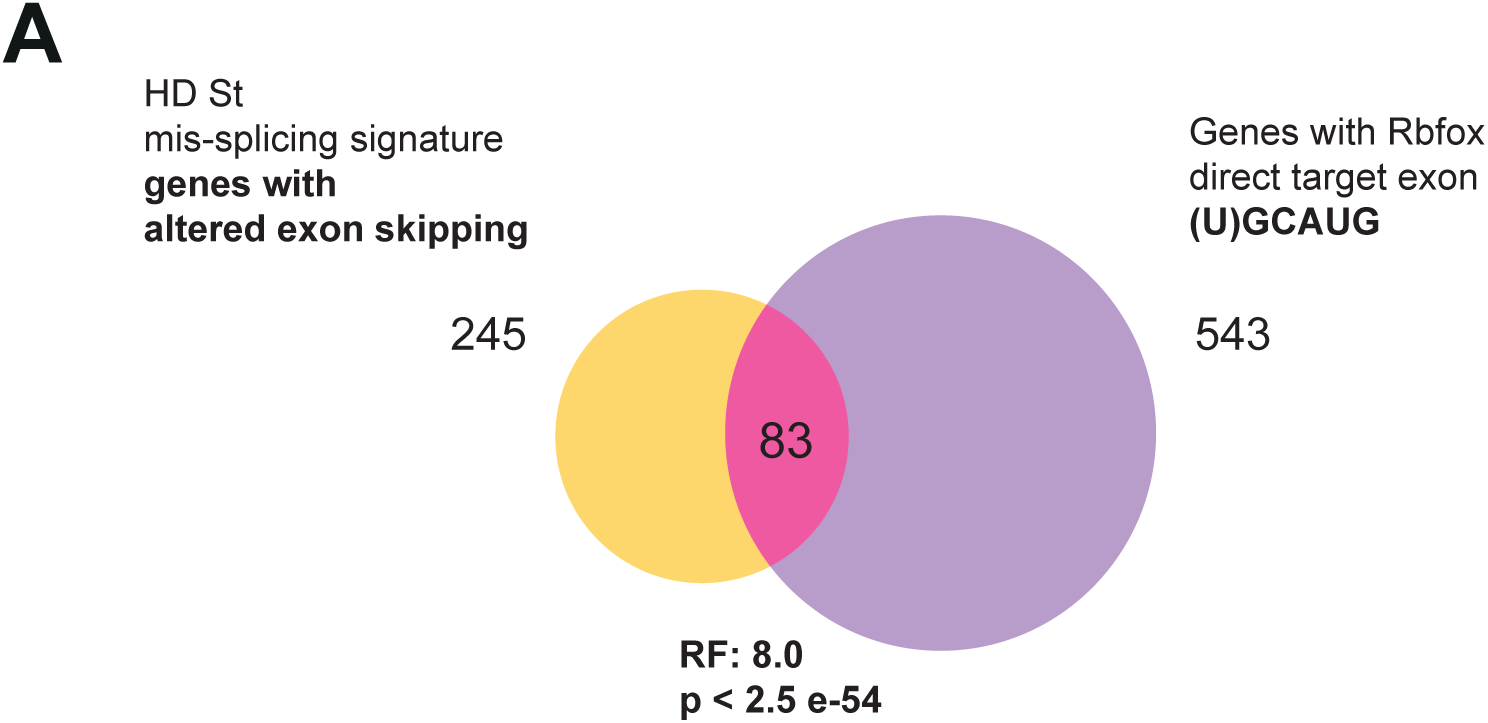
Enrichment of RBFOX direct-targets and Rbfox1 functional targets among Huntington’s disease striatal mis-splicing signature. (**A**) Venn diagram showing the 83 genes in the intersection between the 245 genes with exons mis-spliced in Huntington’s disease^15^ and the 543 genes with RBFOX-direct target exons^25^ (**B**) Venn diagram showing the 76 genes in the intersection between the 245 genes with exons mis-spliced in Huntington’s disease^15^ and the 966 genes that are functional targets of Rbfox1 according to Supplementary Table 1. Representation factor (RF) was determined with Two-sided Fisher’s Exact test, using as background genes the human-mouse orthologous genes coincidentally detected in the human and mouse RNA-seq datasets used to define the Huntington’s disease mis-splicing signature^15^ (*n* = 12,882).

